# Single-cell mass spectrometry reveals heterogeneous triterpenic acid accumulation in apple callus-derived cells

**DOI:** 10.1101/2024.12.08.627150

**Authors:** Carmen Laezza, Sarah Heinicke, Jens Wurlitzer, Vincenzo D’Amelia, Lorenzo Caputi, Maria M. Rigano, Sarah E. O’Connor

## Abstract

The use of plant cell cultures for large scale production of natural compounds, although promising, has been hindered by their genetic instability and heterogeneity. Here, we show how single cell mass spectrometry can be used to characterize the natural product profile of a callus culture at a highly resolved level. We identify and quantify triterpenic acids in a population of callus cells derived from Annurca apple (Malus pumila Miller cv Annurca) leaf. The analysis demonstrated that a high degree of metabolic heterogeneity exists in the cell population, with the levels of detected metabolites varying significantly across the callus cells. This metabolic heterogeneity was underpinned by variable expression levels of key biosynthetic genes in the single cells. The application of an abiotic stress, near ultraviolet radiation (NUV), to the callus culture resulted in increased levels of triterpenic acids. Single cell mass spectrometry analysis revealed that after treatment, a larger percentage of callus cells produced detectable amounts of these metabolites, ultimately resulting in a more homogeneous production of the metabolites. Furthermore, it showed that intracellular concentrations of ursolic acid derivatives can reach more than 100 mM. Single cell mass spectrometry analyses provide a starting foundation for understanding the molecular mechanisms responsible for metabolic heterogeneity in plant cell cultures, which could in turn facilitate efforts to improve these cell cultures for commercial purposes.

---

Plant cell cultures are a promising platform for large scale production of many natural products used in the pharmaceutical, cosmetic and food industries (1). However, despite a few isolated success stories, use of plant cultures in industrial processes is still limited. Although yields of certain metabolites can be increased significantly by abiotic elicitation (2), overall, the biotechnological application of plant cell cultures is hindered by inherent heterogeneity and genetic instability (3). Application of single-cell omics technologies to plant cultures (4) could provide valuable information for development and optimization of high yielding cell cultures. We set out to use our recently established platform for single cell mass spectrometry (scMS) (5) to determine whether the levels of commercially important natural products are produced at similar levels across a population of callus-derived cells.

In this study, we developed a callus culture from leaves of the Annurca apple (*Malus pumila* Miller cv Annurca), a species known to accumulate high levels of triterpenic acids, natural products that possess a wide range of pharmacological properties (6). The interest in these compounds has resulted in increasing commercial demand. However, isolation from plant materials is expensive and often not environmentally sustainable (7), prompting us to develop a plant cell culture for production. After initial metabolomic analysis of the developed callus culture to identify triterpenic acids of interest, we elicited the callus using near-UV radiation (365 nm) for 15 days, which substantially increased the yield of triterpenic acids. We then analyzed small populations of individual NUV treated and non-treated cells using our scMS method (5). scMS revealed that triterpenic acid profiles vary substantially among individual cells of elicited and non-elicited callus, providing a highly resolved understanding of how natural product levels vary among individual cells of plant callus.

Based on the exact mass and fragmentation patterns, our metabolomic analysis revealed the presence of six known triterpenic acids in the callus (Table S1). Ursolic acid (UA) along with the downstream derivatives corosolic acid (CA), euscaphic acid (EA) and annurcoic acid (AA) were detected, in addition to oleanolic (OA) and maslinic acid (MA) (Fig. 1A). With the exception of AA, the identity of these molecules was confirmed by comparison with commercial standards and their accurate quantification was performed by UHPLC-tandem mass spectrometry (Table S2). AA was putatively assigned based on exact mass and comparison of MS^2^ fragmentation with previously reported data (Figures S1, S2) (8). The amounts of individual compounds measured in the callus were 2.09 ± 0.21 (UA), 4.54 ± 1.02 (CA), 4.73 ± 0.73 (EA), 0.47 ± 0.06 (OA) and 2.32 ± 0.32 (MA) mg g^-1^ fresh weight (FW). AA was not quantified due to lack of a standard. To explore the possibility of boosting triterpenic acid production in the callus, which is usually grown in the dark, we exposed callus cultures to both light and light supplemented with NUV radiation (365 nm), which are known to act as abiotic elicitors (9). We observed that near-UV radiation increased the amounts of certain triterpenic acids after 15 days of exposure. Most notably, the levels of CA and EA increased by 2.6- and 2.8-fold, respectively, following elicitation (Fig. S3, Table S3).

**Figure 1.**
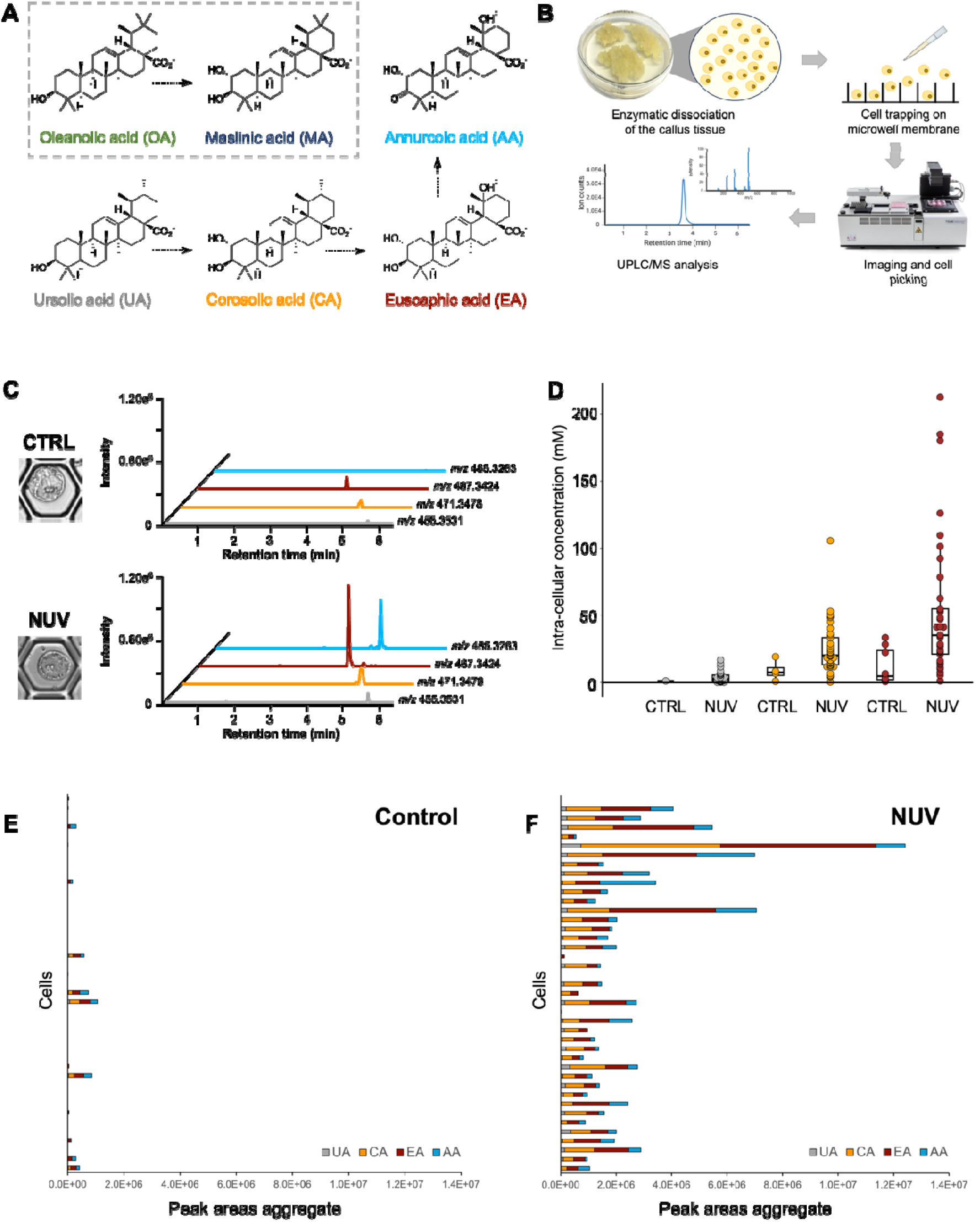
Single cell mass spectrometry (scMS) analysis of callus-derived cells for the detection and quantification of selected triterpenic acids. **A**) The biosynthetic relationships of triterpenic acids detected and quantified in the callus tissue. **B**) Overview of the scMS platform. Cells constituting the callus tissue are enzymatically dissociated (“protoplasting”) and trapped in the 50 μm-wide pores of a microwell membrane. A cell picking robot is then used to image the cells and individually transfer them into 96-well plates. Cells are lysed and further solubilized with organic solvent. The cell extracts are analyzed by UHPLC-HRMS for detection and quantification of the metabolites of interest. **C**) Extracted ion chromatograms of UA (*m/z* 455.3531), CA (*m/z* 471.3478), EA (*m/z* 487.3424), AA (*m/z* 485.3263) in control (CTRL) and near-UV (NUV) treated cells. **D**) Comparison of the intra-cellular concentrations of UA, CA and EA in control and near-UV treated single cells. **E**) Aggregate peak areas of UA, CA, EA and AA measured in control cells. **F**) Aggregate peak areas of UA, CA, EA and AA measured in near-UV treated. All traces, graphs and compounds are color-coded: UA in grey, CA in orange, EA in brown and AA in light blue.

We then applied our scMS method (5) to assess the heterogeneity of triterpenic acid accumulation in the cells constituting the callus, as well as to assess the effect of near-UV treatment on individual cells (Fig. 1B). After enzymatic dissociation of the callus to release individual cells (protoplasts) approximately 10,000 protoplasts were distributed and trapped onto a microwell membrane with 50 μm micropore size and imaged by bright-field microscopy to record size and morphology. Single cells were picked using a microfluidics-based robot and transferred to a 96-well plate (Fig. 1B). After lysis by osmotic shock and solubilization of the released metabolites by addition of methanol, the resulting single cell extracts were measured in negative ionization mode. A set of 40 cells from callus grown in the dark and 40 cells treated with near-UV was successfully analyzed. Extracted ion chromatograms of the targeted analytes from control and near-UV treated cells showed that our analytical method was suitable for the detection of triterpenic acids in single cells (Fig. 1C, Fig. S4). Chromatographic separation of UA and OA, and also CA and MA, which are structural isomers, proved challenging with our single cell setup. Thus, quantification of UA and CA was based on combined peak areas. Previous studies have shown that the short process of releasing the protoplasts (2.5 h) does not impact significantly the levels of natural products (10); however, we cannot exclude that small amounts of metabolites leaked from the cells during the process. With these caveats, we were able to quantify the intracellular concentrations of UA, CA and EA in single cells (Fig. 1C, 1D, Tables S4, S5). UA, CA and EA were detected only in a small number of cells derived from the control callus grown in the dark, but in the few cells that did contain these compounds, the intracellular concentrations were substantial (between 0.69 and 33.43 mM). CA and EA were the most abundant compounds, consistent with the bulk analysis of the entire callus. Elicitation with near-UV light resulted in increased accumulation of triterpenic acids in individual cells (Fig. 1C-F), reaching levels above 200 mM in the case of EA and 100 mM in the case of CA. The data distribution of the measured concentrations of CA and EA was broad, but the median values were 20.19 and 35.19 mM, respectively (Fig. 1D). Hence, production of CA and EA seemed to be favourably affected by NUV-treatment, hinting to an increased rate of hydroxylation of the precursor, UA, by biosynthetic enzymes that have not yet been identified (Fig. 1A). Remarkably, the near-UV elicitation also impacted the number of cells that accumulated the targeted triterpenic acids. The limit of detection was estimated to be an intracellular concentration of approximately 0.1 mM, assuming average cell volume of 13 pL.

AA is a distinctive compound of Annurca apple and a major contributor to the diversity of triterpenic acids in the leaf-derived callus. Since we could not include this metabolite in our quantification study because of the lack of a reference compound, we evaluated the aggregate peak areas of the targeted metabolites detected in the cells (Fig. 1E, 1F). AA production was also affected by near-UV radiation, which again points to stimulation of UA biosynthesis and of downstream oxidative enzyme activities (Fig. 1A). The effect of near-UV elicitation increased the homogeneity of the cell population in terms of triterpenic acid level; specifically, a larger proportion of cells produced detectable levels of triterpenic acids (Fig. 1E, 1F). To corroborate this finding, we performed additional analysis of a larger number (96 per group) of treated and non-treated cells (Fig. S5, S6), which showed a similar pattern. Overall, the control cells produced limited amounts of triterpenic acids, as per their aggregate peak areas, and these were only detectable in approximately 20% of the cells. Following elicitation, the number of cells in which these compounds accumulated increased to 70%; however, a great variability in the quantity of metabolites and their relative ratios still existed.

In conclusion, we used a scMS method to analyze cells of a callus tissue that produces significant amounts of pharmaceutically important triterpenic acids. We measured the intracellular concentrations of three triterpenic acids of interest in a population of these cells. We showed that individual callus cells accumulated variable amounts of the target compounds, revealing the extent of metabolic heterogeneity within the callus cell population. Abiotic elicitation, specifically NUV, dramatically increased the levels of triterpenic compounds. Although the majority of NUV elicited cells sampled contained triterpenic acids, the ratio of the specific triterpenic acids varied substantially across the cell population. Single cell mass spectrometry analyses provide a starting foundation for understanding the molecular mechanisms responsible for metabolic heterogeneity in plant cell cultures, which could in turn facilitate efforts to improve these cell cultures for commercial purposes.

## Acknowledgements

CL was funded by the Erasmus + KA1 Istruzione superiore/Mobilità per l’apprendimento n. 2021-1-IT02-KA131-HED-000011202 e 2022-1-IT02-KA131-HED-000055839 by SEND Consortium. S.E.O. acknowledges the Max Planck Society and the National Institutes of Health (R01 AT012783-02). We thank Anh H. Vu and Mohammed O. Kamileen for helpful discussions.

## Data availability

## Supporting information

Supporting Information file containing supplementary Figures, Tables, Methods and References.

