## Supplementary information for "Single-cell mass spectrometry reveals heterogeneous triterpenic acid accumulation in apple callus-derived cells"

**Author Contributions:** C.L. and L.C. designed and performed the experiments. S.H. assisted with UHPLC-MS method development; J.W. helped with preparing and picking the cells. V.D'A. provided support with the development of callus cultures and with paper revisions; C.L., L.C., M.M.R. and S.E.O'C. conceptualized the study and wrote the paper.

**Competing Interest Statement:** No competing interests to declare.

**Keywords:** single cell mass spectrometry, near-UV elicitation, callus culture, Annurca apple, triterpenic acids.

### This file includes:

**Figures S1 to S6**

**Tables S1 to S5**

**Methods**

**Supplementary references**

### Supplementary Figures

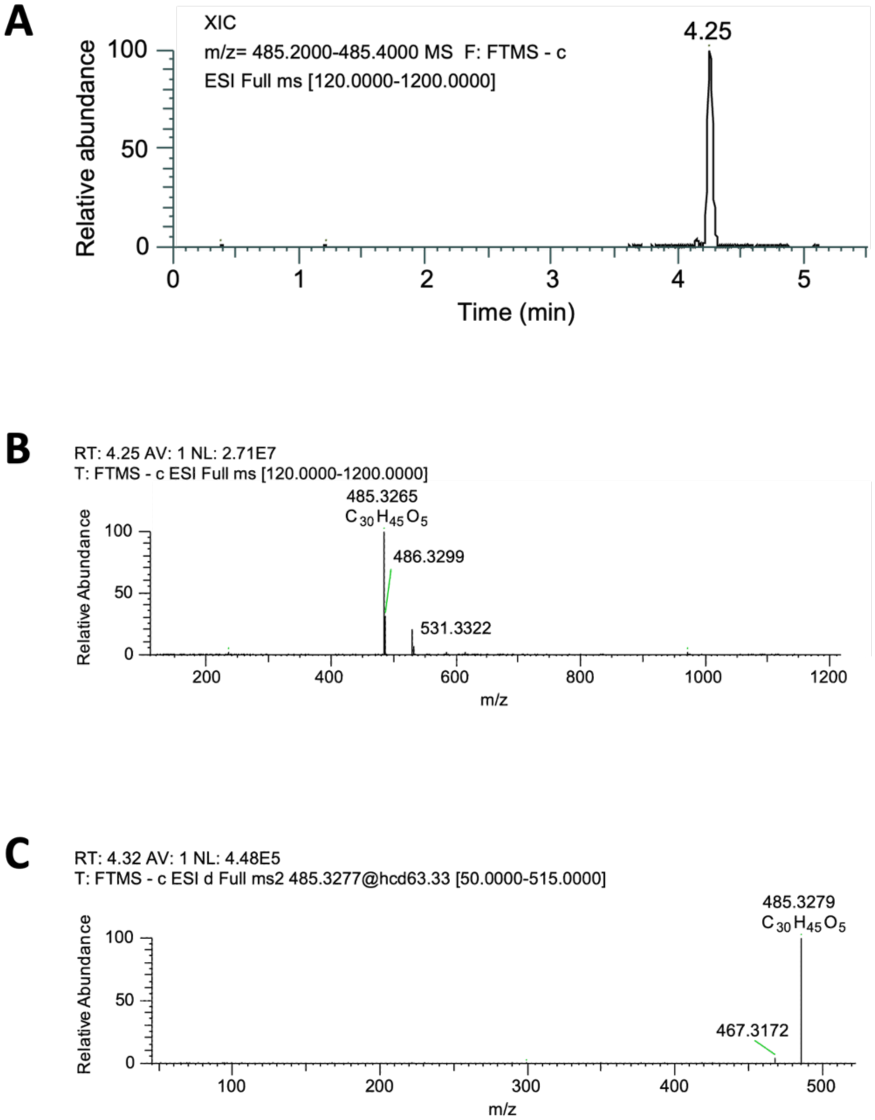


**Figure S1.** High resolution mass spectrometric characterization in negative ionization mode of the putative annurcoic acid peak detected in leaf-derived callus culture. **A**) Extracted ion chromatogram (XIC) of *m/z* 485.20-485.400. **B**) MS chromatogram of the peak at retention time 4.25 min. **C**) MS/MS spectrum of the compound. A fragment at *m/z* 467.3172 $[M\text{-}H\text{-}H_2O]^-$ was observed at the average HCD of 63.33.

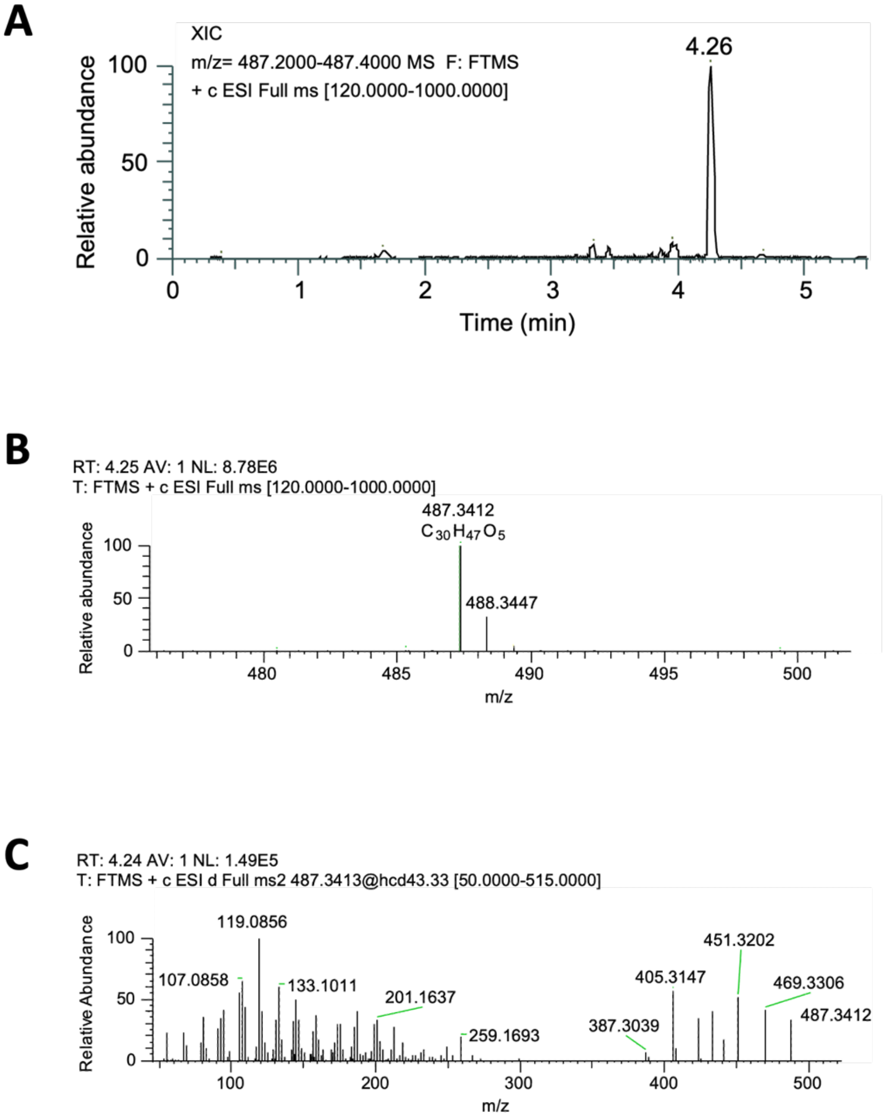


**Figure S2.** High resolution mass spectrometric characterization in positive ionization mode of the putative annurcoic acid peak detected in leaf-derived callus culture. **A**) Extracted ion chromatogram (XIC) of *m/z* 487.20-487.400. **B**) MS chromatogram of the peak at retention time 4.25 min. **C**) MS/MS spectrum of the compound. In positive ionization mode it was possible to fragment the compound.

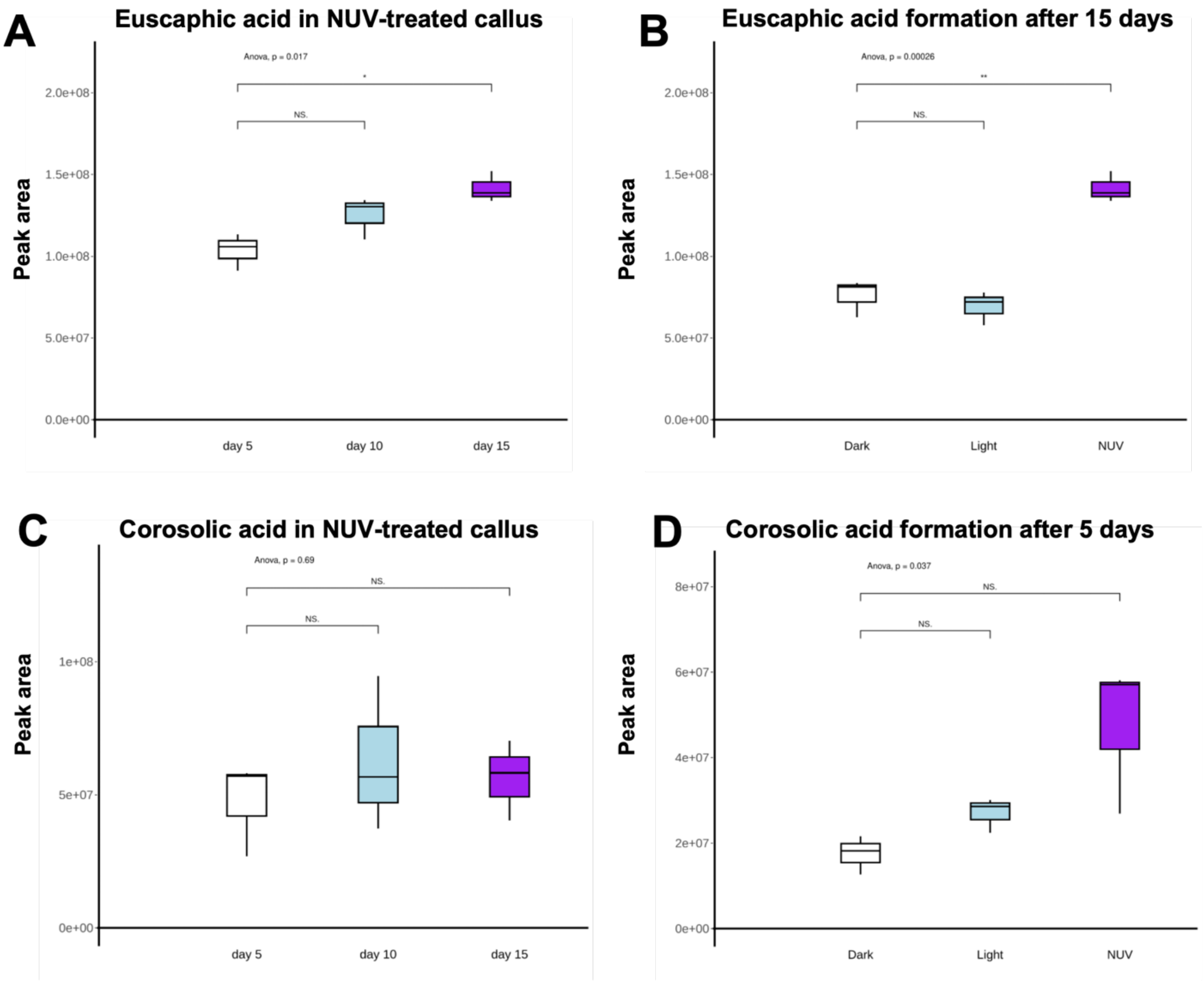


**Figure S3.** Variation in the peak areas of two triterpenic acids during near-UV elicitation as determined by bulk metabolomic analysis of the callus tissue. **A**) During near-UV treatment, the peak area of euscaphic acid increased significantly over time. **B**) After 15 days, the peak area of euscaphic acid was significantly higher in callus treated with near-UV samples compared to those grown in normal light and in the dark. **C**) During near-UV treatment, the peak area of corosolic acid did not increase significantly after 5 days of treatment, likely because it is converted to EA (see panel D and Fig.1A in the main text for the biosynthetic relationship between CA and EA). **D**) After only 5 days, the peak area of corosolic acid was significantly higher in callus treated with near-UV samples compared to those grown in normal light and in the dark.

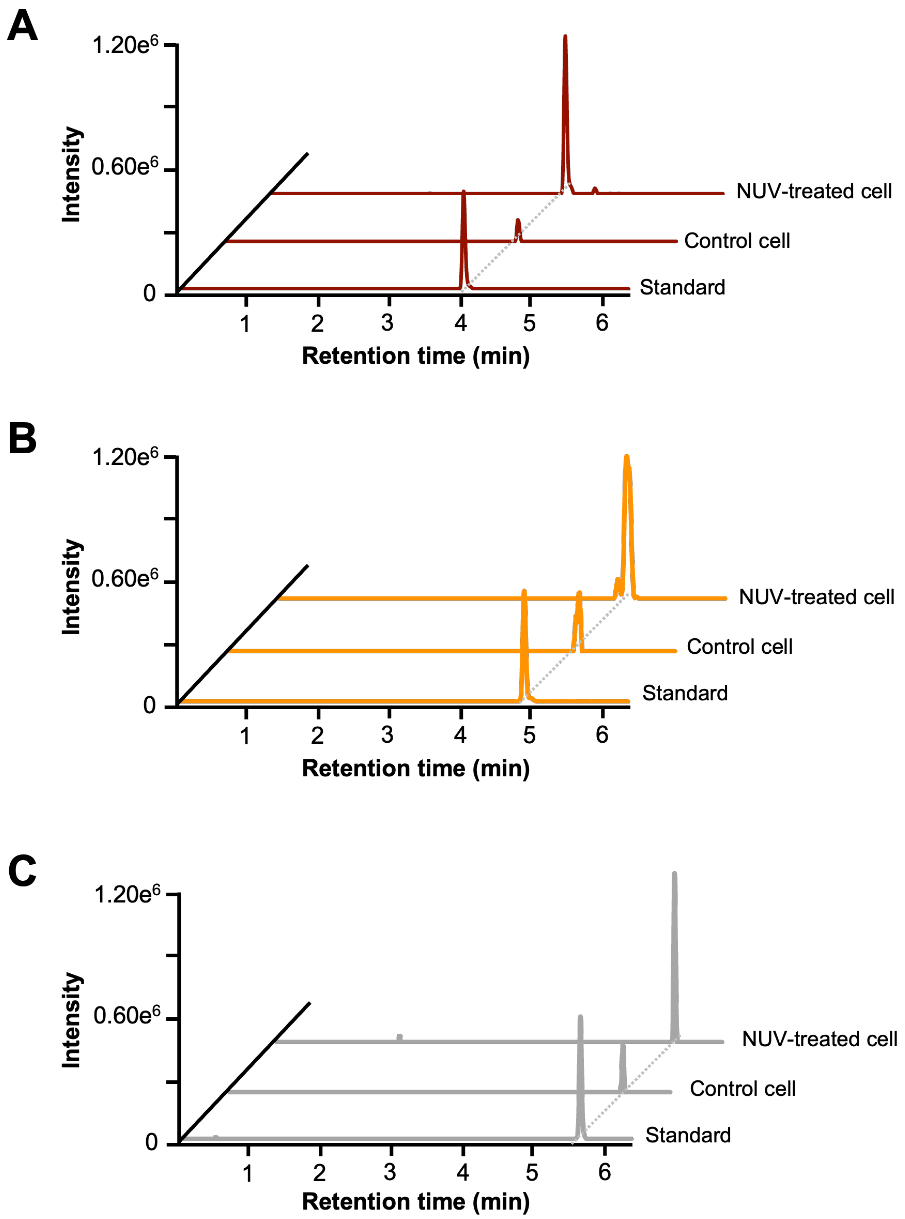


**Figure S4. A**) Comparison of the extracted ion chromatograms of euscaphic acid (*m/z* 487.20-487.40) observed in a control cell and a near-UV treated single cell, in comparison to the authentic standard. **B**) Comparison of the extracted ion chromatograms of corosolic acid (*m/z* 471.20-471.40) observed in a control cell and a near-UV treated cell, in comparison to the authentic standard. The peak shape in the chromatograms from the cells is not perfectly symmetrical, as CA co-elutes with MA. **C**) Comparison of the extracted ion chromatograms of ursolic acid (*m/z* 455.20-455.40) observed in a control cell and a near-UV treated cell, in comparison to the authentic standard. In these chromatographic conditions, UA cannot be separated from OA.

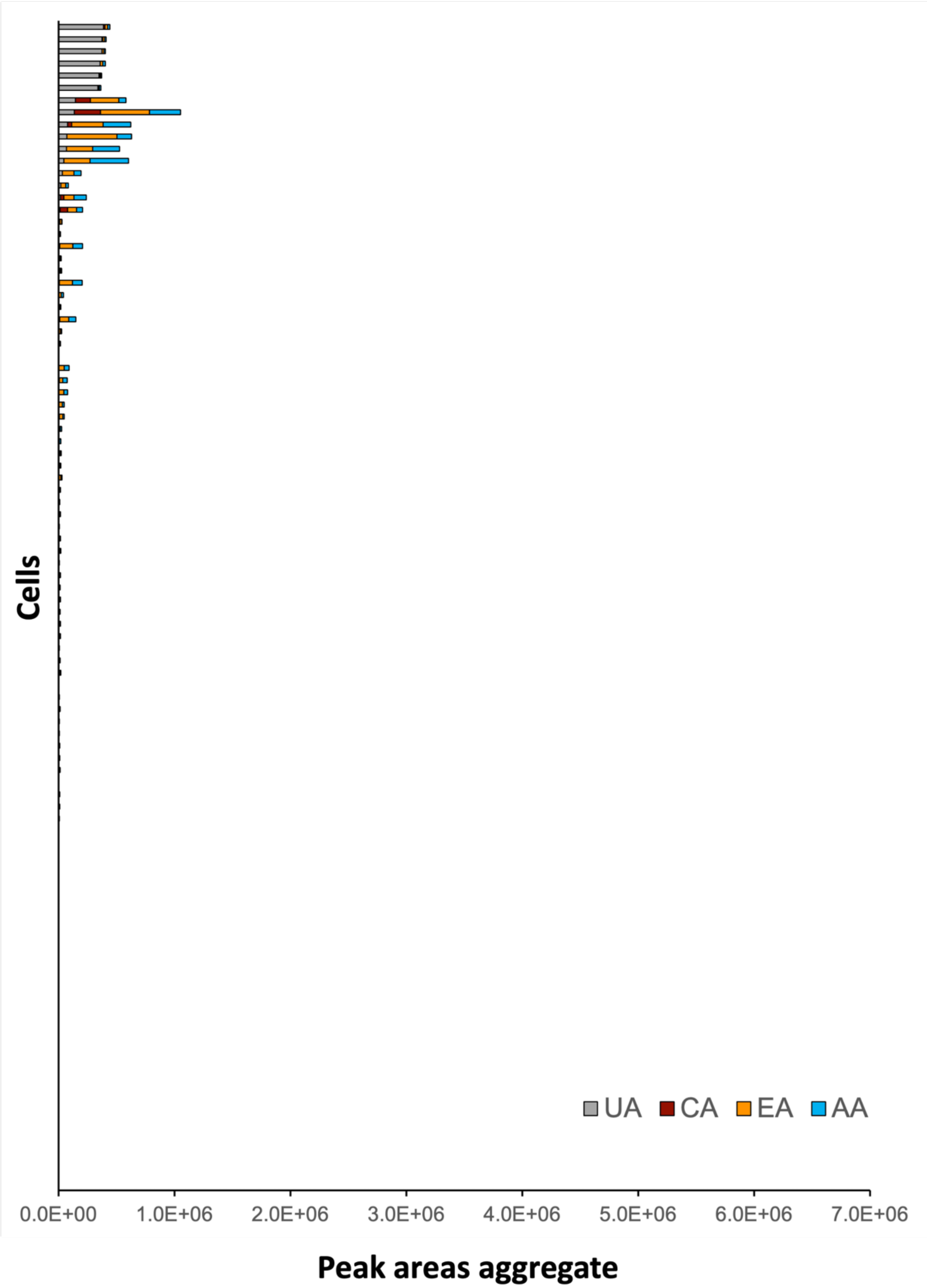


**Figure S5.** Aggregate peak areas of UA, CA, EA and AA measured in control cells. Each bar graph represents a cell.

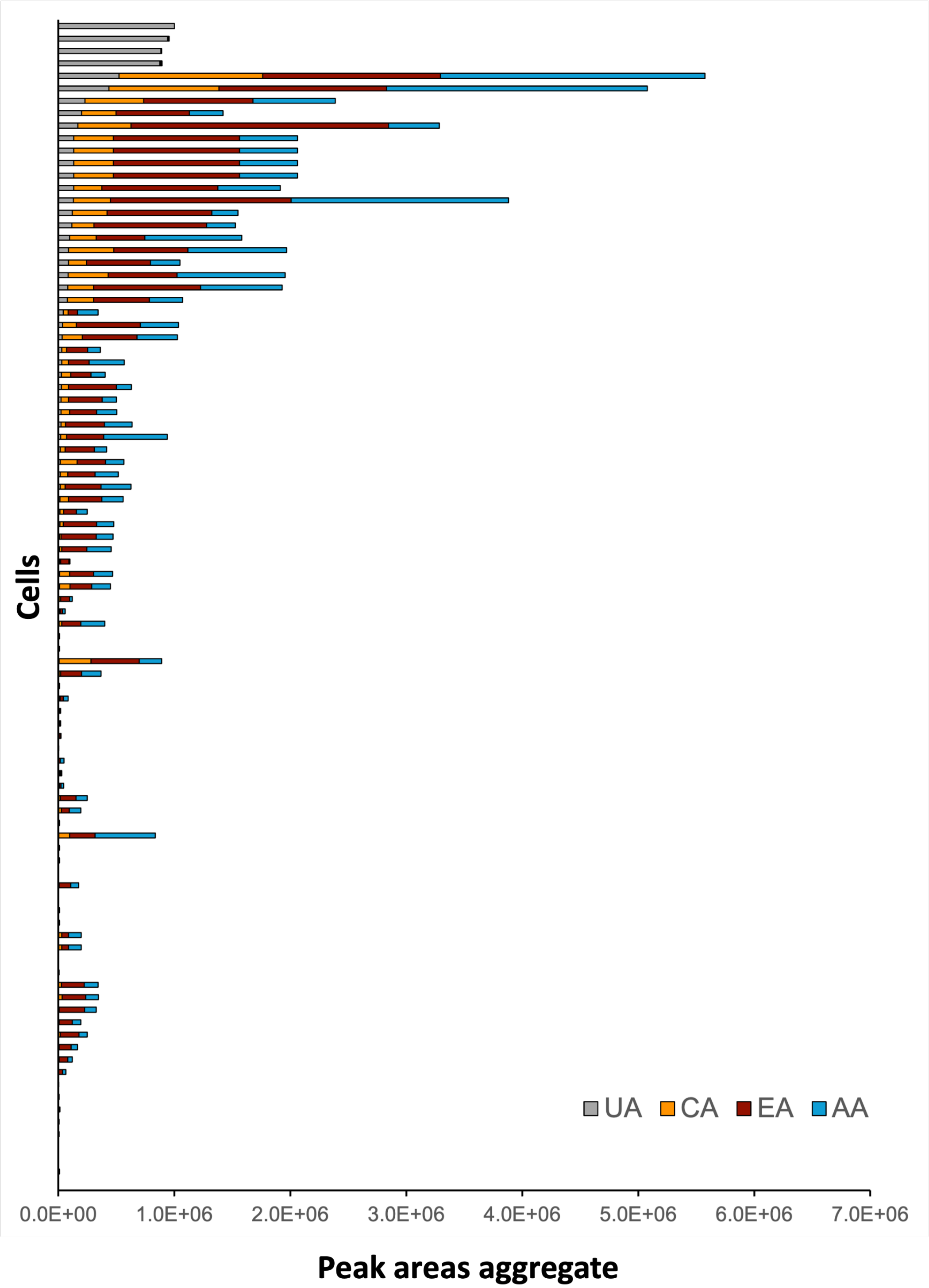


**Figure S6.** Aggregate peak areas of UA, CA, EA and AA measured in near-UV treated cells. Each bar graph represents a cell.

### Supplementary tables

**Table S1.** Triterpenic acids detected by UHPLC-HRMS in Annurca leaf callus [a]

| Molecular formula | Ion mass $[M - H]^-$ | RT | Zodiac score (%) | Sirius score (%) | Tree score | Isotope score | Number of explained peaks | Mass error precursor (ppm) | Predicted structure |
|---|---|---|---|---|---|---|---|---|---|
| $C_{30}H_{44}O_5$ | 483.3178 | 4.49 | 100.00 | 99.98 | 25.90 | 6.06 | 5/9 | 0.94 | COCONUT |
| $C_{30}H_{44}O_6$ | 499.3077 | 3.74 | 100.00 | 100.00 | 28.78 | 6.41 | 5/8 | 0.67 | COCONUT |
| $C_{30}H_{46}O_4$ | 469.3332 | 4.54 | 99.45 | 98.72 | 23.77 | 0 | 4/6 | 0.99 | ChEBI |
| $C_{30}H_{46}O_4$ | 469.3331 | 5.06 | 95.68 | 95.58 | 3.47 | 8.64 | 0/0 | 0.32 | - |
| $C_{30}H_{46}O_5$ | 485.3328 | 4.16 | 100.00 | 99.98 | 50.41 | 3.48 | 11/19 | 0.97 | ChEBI |
| $C_{30}H_{46}O_6$ | 501.3228 | 3.35 | 92.28 | 91.65 | 12.50 | 0 | 1/2 | 1.73 | - |
| $C_{30}H_{46}O_6$ | 501.3227 | 3.54 | 100.00 | 100.00 | 36.71 | 8.36 | 7/9 | 1.81 | COCONUT |
| $C_{30}H_{46}O_7$ | 517.3228 | 3.62 | 94.97 | 95.59 | 4.43 | 6.49 | 1/1 | 1.53 | - |
| $C_{30}H_{48}O_3$ | 455.3545 | 5.25 | 99.40 | 99.50 | 3.38 | 6.47 | 0/0 | 1.38 | - |
| $C_{30}H_{48}O_4$ | 471.3485 | 4.36 | 100.00 | 99.98 | 12.68 | 6.38 | 4/6 | 1.30 | ChEBI |
| $C_{30}H_{48}O_4$ | 471.3486 | 4.47 | 99.95 | 99.98 | 7.71 | 8.46 | 1/5 | 0.46 | - |
| $C_{30}H_{48}O_4$ | 471.3486 | 4.58 | 41.26 | 41.70 | 7.08 | 0 | 2/8 | 0.24 | ChEBI |
| $C_{30}H_{48}O_5$ | 487.3436 | 3.87 | 100.00 | 100.00 | 39.65 | 8.85 | 10/20 | 0.65 | ChEBI |
| $C_{30}H_{48}O_5$ | 487.3441 | 4.19 | 73.92 | 74.82 | 3.35 | 2.32 | 0/0 | 1.75 | - |
| $C_{30}H_{50}O_4$ | 473.3646 | 4.05 | 100.00 | 99.98 | 15.72 | 8.76 | 3/7 | 0.96 | COCONUT |
| $C_{30}H_{50}O_5$ | 489.3590 | 3.44 | 99.40 | 99.50 | 7.16 | 6.54 | 1/4 | 1.52 | - |
| $C_{30}H_{50}O_5$ | 489.3589 | 3.94 | 90.19 | 89.43 | 3.28 | 6.22 | 0/0 | 2.47 | |
| $C_{30}H_{50}O_6$ | 505.3545 | 3.92 | 91.95 | 91.72 | 5.23 | 6.21 | 3/4 | 1.80 | COCONUT |

[a] The table contains: molecular formula; ion mass in negative ionization mode; retention time (RT); Zodiac score; Sirius score; tree score, indicating the computed fragmentation tree; isotope score, indicating the similarity score comparing the measured isotope pattern with the theoretical pattern for each candidate molecular formula; number of explained peaks The number of peaks in the spectrum which can be explained by the fragmentation tree; mass error precursor, indicating the allowed mass deviation in ppm; predicted structure, indicating the databanks in which the molecular structure is already annotated.

**Table S2.** Compounds quantified in the leaf-derived callus tissue by UHPLC-QqQ and parameters of the external calibration using reference compounds.

| Compound name | Calibration range (ng/mL) | Regression equation | Correlation coefficient ($R^2$) |
|---|---|---|---|
| Ursolic acid | 10-300* | y=1670x+2388 | 1.00 |
| Corosolic acid | 10-400 | y=1334x+5698 | 0.99 |
| Euscaphic acid | 10-400 | y=2834x+9763 | 0.99 |
| Oleanolic acid | 10-400 | y=1837x+2445 | 1.00 |
| Maslinic acid | 10-400 | y=1861x+10000 | 0.99 |

*The detector response above 300 ng/mL was not linear

**Table S3**. Concentration (mg $g^{-1}$ FW) of targeted triterpenic acids in control (CTRL), light- and near-UV treated calli [a]

| Metabolites | CTRL | Light-treated | NUV-treated |
|---|---|---|---|
| UA | 2.26 ± 0.30 | 2.95 ± 0.49 | 3.96 ± 0.91* |
| CA | 6.37 ± 2.01 | 5.44 ± 2.20 | 16.53 ± 0.19** |
| EA | 6.13 ± 1.90 | 5.68 ± 1.85 | 17.54 ± 0.50*** |
| OA | 0.73 ± 0.11 | 0.65 ± 0.13 | 1.17 ± 0.23* |
| MA | 4.33 ± 0.70 | 5.04 ± 0.21 | 7.61 ± 0.70** |

[a] Data shown are means ± S.D. Asterisks show statistically significant differences of NUV treatment compared to the control samples (* $p<0.1$; ** $p<0.01$; *** $p<0.001$) by Student's *t*-test.

**Table S4.** Compounds quantified in single cells using the scMS method and parameters of the external calibration using pure analytical standards.

| Compound name | Calibration range (nM) | Regression equation | Correlation Coefficient ($R^2$) |
|---|---|---|---|
| Ursolic acid | 1.95-62.50 | y = 26262x + 52395 | 0.99 |
| Corosolic acid | 1.95-250.00 | y = 30824x + 43191 | 1.00 |
| Euscaphic acid | 1.95-125.00 | y = 45415x + 19621 | 1.00 |

**Table S5.** Intra-cellular quantification of selected triterpenic acids using the scMS method.

| Cell number | Cell diameter (μm) | Cell volume (pL) | Ursolic acid (mM) | Corosolic acid (mM) | Euscaphic acid (mM) |
|---|---|---|---|---|---|
| CTRL_01 | 25.63 | 8.82 | - | - | 6.59 |
| CTRL_02 | 27.59 | 11.00 | - | - | - |
| CTRL_03 | 26.39 | 9.62 | - | - | 2.15 |
| CTRL_04 | 27.24 | 10.58 | - | - | - |
| CTRL_05 | 24.52 | 7.72 | - | - | - |
| CTRL_06 | 32.60 | 18.14 | - | - | - |
| CTRL_07 | 25.06 | 8.24 | - | - | - |
| CTRL_08 | 20.52 | 4.52 | - | - | - |
| CTRL_09 | 37.64 | 27.92 | - | - | - |
| CTRL_10 | 23.31 | 6.63 | - | 8.34 | 28.41 |
| CTRL_11 | 20.59 | 4.57 | - | - | - |
| CTRL_12 | 26.97 | 10.27 | - | - | - |
| CTRL_13 | 28.78 | 12.48 | - | - | - |
| CTRL_14 | 20.41 | 4.45 | - | | - |
| CTRL_15 | 34.41 | 21.33 | - | - | - |
| CTRL_16 | 27.82 | 11.27 | - | - | - |

| Cell number | Cell diameter (μm) | Cell volume (pL) | Ursolic acid (mM) | Corosolic acid (mM) | Euscaphic acid (mM) |
|---|---|---|---|---|---|
| CTRL_17 | 31.32 | 16.09 | - | - | - |
| CTRL_18 | 22.98 | 6.35 | 1.3 | 19.22 | 33.43 |
| CTRL_19 | 22.30 | 5.81 | - | 6.77 | 22.31 |
| CTRL_20 | 31.50 | 16.37 | - | - | - |
| CTRL_21 | 25.12 | 8.30 | - | - | - |
| CTRL_22 | 26.17 | 9.38 | - | - | - |
| CTRL_23 | 44.31 | 45.55 | - | 0.85 | 2.82 |
| CTRL_24 | 33.42 | 19.54 | - | - | - |
| CTRL_25 | 29.84 | 13.91 | - | - | - |
| CTRL_26 | 24.76 | 7.95 | - | - | - |
| CTRL_27 | 31.40 | 16.21 | - | - | - |
| CTRL_28 | 28.21 | 11.75 | - | - | - |
| CTRL_29 | 27.10 | 10.42 | - | - | - |
| CTRL_30 | 28.64 | 12.30 | - | - | - |
| CTRL_31 | 29.44 | 13.36 | - | - | 1.74 |
| CTRL_32 | 34.82 | 22.10 | - | - | - |
| CTRL_33 | 25.62 | 8.81 | - | - | - |
| CTRL_34 | 34.58 | 21.65 | - | - | - |
| CTRL_35 | 31.57 | 16.47 | - | - | - |
| CTRL_36 | 23.55 | 6.84 | - | - | - |
| CTRL_37 | 31.47 | 16.32 | - | - | 0.69 |
| CTRL_38 | 32.88 | 18.61 | - | - | - |
| CTRL_39 | 32.65 | 18.22 | - | - | - |
| CTRL_40 | 39.65 | 32.64 | - | - | - |
| NUV_01 | 26.61 | 9.87 | - | 5.30 | 22.72 |
| NUV_02 | 38.58 | 30.07 | 0.58 | 4.27 | 7.50 |
| NUV_03 | 29.81 | 13.87 | 2.86 | 28.90 | 52.73 |
| NUV_04 | 20.23 | 4.33 | 2.08 | 33.37 | 126.02 |

| Cell number | Cell diameter (μm) | Cell volume (pL) | Ursolic acid (mM) | Corosolic acid (mM) | Euscaphic acid (mM) |
|---|---|---|---|---|---|
| NUV_05 | 28.50 | 12.12 | 11.81 | 21.88 | 28.46 |
| NUV_06 | 26.57 | 9.82 | - | 6.73 | 23.89 |
| NUV_07 | 27.90 | 11.37 | 3.68 | 25.30 | 21.12 |
| NUV_08 | 26.54 | 9.79 | - | 14.91 | 78.47 |
| NUV_09 | 27.34 | 10.70 | 1.82 | 11.81 | 16.18 |
| NUV_10 | 22.19 | 5.72 | 1.99 | 26.91 | 41.22 |
| NUV_11 | 42.72 | 40.82 | 3.12 | 11.91 | 11.24 |
| NUV_12 | 21.91 | 5.51 | 0.84 | 22.69 | 23.72 |
| NUV_13 | 20.52 | 4.52 | 14.06 | 52.97 | 47.18 |
| NUV_14 | 28.04 | 11.54 | 0.32 | 12.76 | 28.58 |
| NUV_15 | 27.10 | 10.42 | 3.43 | 18.48 | 13.52 |
| NUV_16 | 30.57 | 14.96 | 0.97 | 14.60 | 41.18 |
| NUV_17 | 27.59 | 11.00 | - | - | - |
| NUV_18 | 23.71 | 6.98 | 5.85 | 48.51 | 109.62 |
| NUV_19 | 25.71 | 8.90 | - | 11.80 | 14.45 |
| NUV_20 | 32.93 | 18.70 | 1.53 | 13.23 | 15.96 |
| NUV_21 | 28.36 | 11.94 | - | - | - |
| NUV_22 | 32.51 | 17.99 | 2.54 | 16.64 | 10.23 |
| NUV_23 | 29.41 | 13.32 | - | 0.37 | 1.17 |
| NUV_24 | 28.66 | 12.33 | 3.67 | 22.42 | 27.12 |
| NUV_25 | 27.06 | 10.37 | 1.50 | 19.76 | 35.39 |
| NUV_26 | 23.93 | 7.18 | 6.97 | 50.27 | 48.36 |
| NUV_27 | 29.21 | 13.05 | 0.23 | 20.19 | 41.53 |
| NUV_28 | 29.03 | 12.81 | 6.41 | 45.07 | 180.11 |
| NUV_29 | 24.28 | 7.49 | 2.39 | 18.51 | 33.78 |
| NUV_30 | 30.93 | 15.49 | 1.39 | 16.37 | 23.12 |
| NUV_31 | 24.87 | 8.05 | 1.18 | 20.41 | 62.58 |
| NUV_32 | 24.74 | 7.93 | 4.00 | 40.31 | 92.37 |

| Cell number | Cell diameter (μm) | Cell volume (pL) | Ursolic acid (mM) | Corosolic acid (mM) | Euscaphic acid (mM) |
|---|---|---|---|---|---|
| NUV_33 | 28.69 | 12.36 | 1.82 | 14.35 | 35.19 |
| NUV_34 | 26.40 | 9.63 | 8.58 | 49.51 | 212.42 |
| NUV_35 | 32.76 | 18.41 | 16.78 | 105.58 | 184.83 |
| NUV_36 | 30.86 | 15.39 | 0.21 | 4.58 | 5.33 |
| NUV_37 | 32.00 | 17.16 | 5.46 | 36.34 | 101.36 |
| NUV_38 | 27.41 | 10.78 | 7.24 | 35.33 | 54.86 |
| NUV_39 | 33.47 | 19.63 | 3.37 | 24.17 | 54.15 |
| NUV_40 | 22.82 | 6.22 | - | - | - |

### Methods

#### Chemicals and reagents

Ursolic acid and oleanolic acid were purchased from TCI (Tokyo Chemical Industry Co.). Maslinic acid was obtained from Merk, whilst corosolic acid and euscaphic acid were obtained from Toronto Research Chemicals Inc. Milli-Q water was used to prepare all solutions. Murashige and Skoog (MS) medium, plant growth regulators (2,4-dichlorophenoxyacetic acid and 6-benzylaminopurine) and sucrose were purchased from Duchefa Biochemie (RV Haarlem, Netherlands). For protoplast extraction, Cellulase Onozuka R-10, Macerozyme R-10 were from SERVA, whilst pectinase, mannitol, KCl and MES were purchased from Merk. $CaCl_2 \times 2H_2O$ was from Carl Roth. All solvents used in this study were of UHPLC/MS grade.

#### Plant material and establishment of the callus culture

Callus cultures were developed from the leaves of the apple Malus pumila Miller cv Annurca. The sterilization of leaf tissue was performed as described by Laezza *et al*. (1). MS medium was prepared by mixing 30 g $L^{-1}$ sucrose, 2.5 mg $L^{-1}$ 2,4-dichlorophenoxyacetic acid (2,4-D), 0.5 mg $L^{-1}$ 6-benzylaminopurine (BAP) and 0.8% agar. After adjusting the pH to 5.8, the medium was autoclaved at 121 °C for 20 min and poured in 90 mm Petri dishes. Small pieces of sterile Annurca leaves were placed on plates containing the medium and incubated in the dark at 25 ± 2 °C. The growing callus was sub-cultured every 15 days.

#### Extraction of triterpenic acids

Calli from exponential developmental stage (10th day) were frozen in liquid nitrogen and ground using mortar and pestle. Extraction of metabolites was performed by addition of 300 µL of 80% methanol containing 0.1% formic acid solution to 10 mg of frozen powder obtained. After mixing vigorously for 1 min, the samples were first placed in an ultrasonic bath for 20 min, then centrifuged at 24000 g for 10 min. The supernatants were collected, filtered through 0.22 µm filters and stored at -20 °C before further analyses.

#### Metabolomic analysis of callus tissue by UHPLC-HRMS

The UHPLC-HRMS analysis was performed on a Vanquish (Thermo Fisher Scientific) system coupled to a Q-Exactive Plus (Thermo Fisher Scientific) orbitrap mass spectrometer. For metabolite separation, a

Waters™ ACQUITY UPLC BEH C18 130 Å column (1.7 µm, 2.1 mm x 50 mm) column was used at a temperature of 40°C. The binary mobile phases were water 0.1 % formic acid (A) and acetonitrile (B). The gradient elution started with 1% ACN and increased linearly to 70% ACN over 5 min. The wash stage was performed at 99% ACN for 0.5 min before switching back to 1% ACN for 1.5 min to condition the column for the next injection. Total time for chromatographic separation was 7 min. The flow rate was 0.6 mL $min^{-1}$ during the chromatographic separation. In total, 2 µL of standards or samples were injected into the column via the autosampler. Both samples and standard solutions were kept at 10 °C in the sample tray. The needle in the autosampler was washed using a mixture of methanol and MilliQ water (1:1, v:v) for 20 s after the draw and at a speed of 50 µL $s^{-1}$. The mass spectrometer was equipped with a heated electrospray ionization source. The mass spectrometer was calibrated using the Pierce positive and negative ion mass calibration solution (Thermo Fisher Scientific). The operating parameters of heated electrospray ionization are based on the UHPLC flow rate of 0.6 mL $min^{-1}$ using source auto default: sheath gas flow rate at 55; auxiliary gas flow rate at 15; sweep gas flow rate at 3; spray voltage +3500 V; capillary temperature at 275 °C; auxiliary gas heater temperature at 450 °C and S-lens RF level at 50.

Acquisition was performed in full-scan MS mode (resolution 70000-FWHM at 200 Da) in negative mode over the mass range *m/z* from 120 to 1000. The full MS/dd-MS2 (full-scan and data-dependent MS/MS mode) was used to simultaneously record the MS/MS (fragmentation) and the spectra for the precursors of QC pooled samples. The full MS/dd-MS2 (that included target analytes) was also used for QC pooled samples to confirm fragments of the selected precursors. The dd-MS2 was set up with the following parameters: resolution 17500 FWHM; mass isolation window 0.7 Da; maximum and minimum automatic gain control target $8 \times 10^3$ and $5 \times 10^3$, respectively; normalized collision energy was set at three levels 10%, 30% and 45% and spectrum data format was centroid. All the parameters of the UPLC-MS system were controlled through Thermo Fisher Scientific Xcalibur software version 4.3.73.11 (Thermo Fisher Scientific). For the characterization of annurcoic acid, spectra were recorded both in positive and negative mode using higher collision energy (30%, 70% and 100%).

**Quantification of triterpenic acids in callus tissue by UHPLC-QqQ**

UHPLC–QqQ analysis of triterpenic acids was performed on a Thermo Scientific UltiMate 3000 RS ultra-high performance liquid chromatography (UHPLC) system (Thermo Scientific) coupled to a EVOQ Elite™ triple quadrupole mass spectrometer (Bruker Daltonics). For metabolite separation, a ZORBAX RRHT Extend C18 column (80 Å, 4.6 x 50 mm, 1.8 µm, Agilent) was used at a temperature of 40 °C. The binary mobile phases were water 0.1 % formic acid (A) and methanol (B). The separation was achieved by isocratic conditions at 10/90 (v/v) of A/B within 5 min of total run time. The injection volume of both the standard solutions and the samples was 2 µL. The flow rate of the mobile phase was kept constant at 0.6 mL $min^{-1}$ during the chromatographic separation. Both samples and standard solutions were kept at

10 °C in the sample tray. The needle in the autosampler was washed using acetonitrile for 5 s before and after draw and at a speed of 20 µL $s^{-1}$. The mass spectrometer was equipped with a heated electrospray ionization source. The mass spectrometer was operated in positive and negative ionization mode simultaneously in one run to achieve the best sensitivity for each analyte of interest. The EVOQ source parameters were as follows: heated ESI spray voltage (+/-) 4000 V; cone gas flow 20 arbitrary units at 350°C; probe gas flow 45 arbitrary units at 450°C; nebulizer gas flow 50 arbitrary units; exhaust gas on. The analysis in negative ionization mode was performed in single ion monitoring (SIM), operating the Q1 mass analyzer under unit resolution (0.7 Da FWHM). The following precursor ions were recorded: *m/z* 471.3 for both maslinic and corosolic acid ($C_{30}H_{48}O_4$), *m/z* 487.1 euscaphic acid ($C_{30}H_{48}O_5$). The analysis in positive ionization was performed in multiple reaction monitoring (MRM) mode operating the Q1 mass analyzer under unit resolution (0.7 Da FWHM) and Q3 mass analyzer at 2.0 Da FWHM. MRM transitions were determined from the analytical standards and used to record during sample analysis: oleanolic acid ($C_{30}H_{48}O_3$) (quantifier: *m/z* 439.0 > 191.1, qualifier 1: *m/z* 439.0 > 203.1, qualifier 2: *m/z* 439.0 > 95.1), ursolic acid ($C_{30}H_{48}O_3$) (quantifier: *m/z* 439.2 > 191.1, qualifier 1: m/z 439.2 > 205.1, qualifier 2: *m/z* 439.2 > 95.1). The identification of each analyte of interest in the respective samples was confirmed by comparing retention times to authentic reference standards which were measured in the same sequence together with the samples. The samples were individually diluted so that they are in the linear range of the calibration curve of the respective analytes. For corresponding stock solutions, ursolic, oleanolic, corosolic, euscaphic and maslinic acids were dissolved in pure MeOH at a concentration of 1 mg $mL^{-1}$. Dilutions were prepared as follows: 10, 25, 30, 50, 80, 100, 200, 300, 400 ng $mL^{-1}$. The respective calibration curves and sample concentrations were calculated using Data Review version 8.2.1 of the MS workstation software (Bruker Daltonics).

**Analysis of triterpenic acids in single cells by scMS**

UHPLC-HRMS analysis was performed on a Vanquish (Thermo Fisher Scientific) system coupled to a Q-Exactive Plus (Thermo Fisher Scientific) orbitrap mass spectrometer. For metabolite separation, a Waters™ ACQUITY UPLC BEH C18 130 Å column (1.7 µm, 1 mm x 50 mm) column was used at a temperature of 40°C. The binary mobile phases were water 0.1 % formic acid (A) and acetonitrile (B).. The gradient elution started with 1% ACN and increased linearly to 70% ACN over 5 min. The wash stage was performed at 99% ACN for 0.5 min before switching back to 1% ACN for 1.5 min to condition the column for the next injection. Total time for chromatographic separation was 7 min. The flow rate was 0.3 mL $min^{-1}$ during the chromatographic separation. In total, 4 µl of standards or samples were injected into the column via the autosampler. Both samples and standard solutions were kept at 10 °C in the sample tray. The needle in the autosampler was washed using a mixture of methanol and MilliQ water (1:1, v:v) for 20 s after the draw and at a speed of 50 µL $s^{-1}$. The mass spectrometer was equipped with a heated electrospray ionization source. The mass spectrometer was calibrated using the Pierce positive and negative ion mass calibration solution (Thermo Fisher Scientific). The operating parameters of heated electrospray ionization are based on the UPLC flow rate of 0.3 mL $min^{-1}$ using source auto default: sheath gas flow rate at 48; auxiliary gas flow rate at 11; sweep gas flow rate at 1; spray voltage +3500 V; capillary temperature at 250 °C; auxiliary gas heater temperature at 300 °C and S-lens RF level at 50.

Acquisition was performed in full-scan MS mode (resolution 70000-FWHM at 200 Da) in negative mode over the mass range *m/z* from 120 to 1,000. The full MS/dd-MS2 (full-scan and data-dependent MS/MS mode) was used to simultaneously record the MS/MS (fragmentation) and the spectra for the precursors of QC pooled samples. The full MS/dd-MS2 (that included target analytes) was also used for QC pooled sample to confirm fragments of the selected precursors. The dd-MS2 was set up with the following parameters: resolution 17,500 FWHM; mass isolation window 0.7 Da; maximum and minimum automatic gain control target $8 \times 10^3$ and $5 \times 10^3$, respectively; normalized collision energy was set at three levels 10%, 30% and 45% and spectrum data format was centroid. All the parameters of the UPLC-HRMS system were controlled through Thermo Fisher Scientific Xcalibur software version 4.3.73.11 (Thermo Fisher Scientific). Chromatography and MS responses were optimized using several reference compounds. To quantify TAs in single cells, calibration curves were prepared. UA, EA and CA were dissolved in pure MeOH at a concentration of 1 mg $L^{-1}$. Serial dilutions (n = 10) were made between 250 nM and 1.95 nM and analyzed by UHPLC–HRMS. The extracted peak areas were used to calculate linear calibration curves.

#### Callus cells dissociation

Light-treated and CTRL leaf-derived calli were used for extracting protoplasts with the purpose of establishing how the triterpenic acid pathway might change at cellular level. First, the digestion medium was prepared; (1.5% (wt/vol) cellulose Onozuka R-10, 0.4% (wt/vol) macerozyme R-10 and 0.1% (vol/vol) pectinase were dissolved in mannitol/MES (MM) buffer. MM buffer contains 400 mM mannitol, 60 mM sucrose, 20 mM KCl, 10 mM CaCl2 x $2H_2O$, and 20 mM MES. The pH of the buffer solution was adjusted to 5.7–5.8 with 1 M KOH. Approximately 0.5 g of callus was added to 4 ml of digestion medium, and the callus was distributed into small plates 40 mm diameter to maximize surface contact with digestion solution with 100 µL pipette tip. Thereafter, the plate containing the callus was incubated for 2 h at room temperature, in the dark, as the enzymes are light-sensitive. After the incubation, the plates were placed on an orbital shaker for 30 min at 70 rpm to release the protoplasts. The protoplast suspension was filtered through a 100 µm cell strainer to remove larger debris and gently transferred to two 15 mL Falcon tubes. The protoplast suspension was centrifuged at 100 g with gentle acceleration/deceleration, for 5 min, at 12 °C to pellet the protoplasts. The supernatant was removed as much as possible, sucking it off with vacuum; these steps were repeated three times to wash the protoplast pellet. Eventually, the residual pellet from the two tubes were pooled together and resuspended in 1 mL of MM buffer. The protoplast concentration was determined using a cell counter LUNA II (Logos Biosystems, Aligned Genetics, Inc.). 20 μL of protoplast cell suspension was mixed with 20 μL of Typane Blue Stain (0.4%) (Gibco). From the mix 10 μL where load in a LUNA™ Cell Counting Slide (Logos Biosystems, Aligned Genetics, Inc.) for measurement. The final concentration of protoplasts was adjusted to 10k protoplasts in 1 mL.

**Cell picking for scMS Analysis**

A SIEVEWELL™ Slide Dimensions (W x D x H): 25 mm x 75 mm x 12 mm (Sartorius) with 90,000 nanowells (50 μm × 50 μm, depth × diameter) was used for single-cell trapping and sorting. The SIEVEWELL chip was primed with 2 mL 100% ethanol to break surface tension. The EtOH was removed from the side ports and replaced by 2 mL of MM buffer in the same way. This washing step was repeated three times. After that, washing the chip was continued with 3 mL (0.5%) BSA in MM-buffer and subsequently discarding the liquid through the side port. The chip's membrane was then allowed to be coated by addition of fresh 0.5% BSA in MM-buffer and incubation O/N at 4°C. Finally, the 0.5% BSA was removed through the side ports and washed away with 2 mL of MM-buffer as described above. Finally, 1 mL pre-diluted protoplast suspension was carefully added and dispensed in a z-shape across the well. Almost all the liquid was then discarded from the side ports leading to sedimentation of the cells on the membrane. The chamber of the slide was filled up with 1mL fresh MM-buffer. The buffer was removed this time from the top of the chamber to remove loose cells and debris and finally replaced by 1mL new MM-buffer. The SIEVEWELL was then mounted on the CellCelector Flex (Sartorius) instrument, and the cells were visualized using the optical unit, constituted by a fluorescence microscope (Spectra X Lumencor) and a CCD camera (XM-10). Images of the cell under transmitted light were acquired during the picking process. Single protoplasts were picked using a 50 μm glass capillary SingleCell 50 (Sartorius). Cells that were used for UHPLC-HRMS analysis were dispensed into SureSTART™ WebSeal™ 96-Well Microtiter Plates, V-Bottom (Thermo Scientific) containing 6 μL of MilliQ water. After the picking, 6 μL of MeOH containing 10 nM of harpagoside (Extrasynthese) as an internal standard were added to improve metabolite solubilization. 2 μL from each well were combined in a UPLC glass vial to constitute the pooled Quality Control (QC) sample. Plates were frozen at −80 °C until analysis.

**Elicitation of the callus culture by near-UV light**

To enhance the production of triterpenic acids, different light treatments were applied to the callus cultures. Calli from exponential developmental stage (10th day) were grown either in the dark (CTRL) or under two light conditions: light, 380-780 nm (LED LightDNA-8 300 W, Valoya LED Grow Lights, Helsinki, Finland) and Near UV light (NUV), in which the LED LightDNA was supplemented with 365 nm radiation generated by a panel of LED UVA Puzzle 5W (Roschwege GmbH, Greifenstein, Germany), at 16 h on/8 h off, 25 ± 2 °C for 15 days. For the metabolomic analysis, the samples were harvested at three time points: after 5 days, 10 days and 15 days.

**Data analysis and visualization**

For metabolomic analysis, mass spectrometry data were converted into mzML files by using MSConvert (ProteoWizard software). These files were analyzed using SIRIUS (v.5.8.5), ZODIAC, CSI:FingerID and CANOPUS to turn mass spectra into structure information of metabolites detected in the samples (2). For targeted analysis, mass spectrometry data were processed using QuanBowser (Bruker Daltonics). UHPLC-MS data were visualized using Freestyle Software (Thermo Fisher Scientific). The diameter of cells was measured with ImageJ (v.1.54).
